# A genome assembly of the barley ‘transformation reference’ cultivar Golden Promise

**DOI:** 10.1101/2020.02.12.945550

**Authors:** Miriam Schreiber, Martin Mascher, Jonathan Wright, Sudharasan Padmarasu, Axel Himmelbach, Darren Heavens, Linda Milne, Bernardo J. Clavijo, Nils Stein, Robbie Waugh

**Author notes:** Corresponding author(s): Robbie Waugh.

## Abstract

**Background:** Barley (*Hordeum vulgare*) is one of the most important crops worldwide and is also considered a research model for the large-genome small grain temperate cereals. Despite genomic resources improving all the time, they are limited for the *cv.* Golden Promise, the most efficient genotype for genetic transformation.

**Findings:** We have developed a barley *cv.* Golden Promise reference assembly integrating Illumina paired-end reads, long mate-pair reads, Dovetail Chicago in vitro proximity ligation libraries and chromosome conformation capture sequencing (Hi-C) libraries into a contiguous reference assembly. The assembled genome of 7 chromosomes and 4.13Gb in size, has a super-scaffold N50 after Chicago libraries of 4.14Mb and contains only 2.2% gaps. Using BUSCO (benchmarking universal single copy orthologous genes) as evaluation the genome assembly contains 95.2% of complete and single copy genes from the plant database.

**Conclusions:** A high-quality Golden Promise reference assembly will be useful and utilised by the whole barley research community but will prove particularly useful for CRISPR-Cas9 experiments.

## Data Description

Barley is a true diploid with 14 chromosomes (2n=14). Its genome is around 5Gb in size and mainly consists of repetitive elements^1^. Barley is and has been an important crop for thousands of years^2^. It was the fourth most produced cereal in 2016 worldwide (Faostat, http://www.fao.org/faostat/en/#home) and second most in the UK. While the majority is used as feed an important market for 2-row spring barley is the whisky industry. An iconic historical variety is *cv.* Golden Promise. Golden Promise is a 2-row spring type which was mainly grown in Scotland in the 1970s and early 1980s and was identified as a semi-dwarf mutant after a gamma-ray treatment of the cultivar Maythorpe. It was used extensively for malting and whisky production and some distilleries still use it today. In recent years, the main research interest in Golden Promise has come from its genetic transformability. While many other cultivars have been tested^3-6^ and some successfully used, the best shoot recovery from callus is usually achieved, and most barley transformations are successfully conducted, using Golden Promise. With the rise of the CRISPR-Cas9 genome editing technology, a potential Golden Promise reference assembly has already sparked wide interest in the barley community. The use of CRISPR-Cas9 ideally requires a complete and correct reference assembly for the identification of target sites. The Cas9 enzyme targets a position in the genome based on a sgRNA (single-guide RNA) followed by a PAM (protospacer-adjacent motif). The guide RNA is usually designed to be 20 bp long and to be specific to the target to avoid any off-target effects. The PAM region consists of three nucleotides “NGG” ^7,8^. Any nucleotide variation between different cultivars can therefore cause problems with the CRISPR-Cas9 genome editing technology. The time and cost involved in such increasingly common experiments highlights the value of a high-quality Golden Promise reference assembly.

## Methods

### Contig construction and scaffolding

#### DNA extraction, library construction and sequencing

High molecular weight barley DNA was isolated from leaf material of 3-week old Golden Promise plants that had been kept in the dark for 48 hours to reduce starch levels. DNA was extracted using the GE Life Sciences Nucleon PhytoPure kit (GE Healthcare Life Sciences, Buckinghamshire, UK) according to the Manufacturers’ instructions. Both paired end and long mate pair libraries were constructed and sequenced at the Earlham Institute by the Genomics Pipelines Group. A total of 2 µg of DNA was sheared targeting 1 kbp fragments on a Covaris-S2 (Covaris Brighton, UK), size selected on a Sage Science Blue Pippin 1.5% cassette (Sage Science, Beverly, USA) to remove DNA molecules <600bp, and amplification-free, paired end libraries constructed using the Kapa Biosciences Hyper Prep Kit (Roche, New Jersey, USA). Long mate pair libraries were constructed from 9 µg of DNA according to the protocol described in Heavens *et al.*, 2015^11^ based on the Illumina Nextera Long Mate Pair Kit (Illumina, San Diego, USA). Sequencing was performed on Illumina HiSeq 2500 instruments with a 2×250 bp read metric targeting >60x raw coverage of the amplification-free library and 30x coverage of a combination of different insert long mate pair libraries with inserts sizes >7 kbp.

#### Contig and scaffold generation

Approximately 624 million 2×250 bp paired reads were generated providing an estimated 62.4x coverage of the genome. Contigging was performed using the w2rap-contigger^12^. Three mate-pair libraries were produced with insert sizes 6.5, 8 and 9.5kb and sequenced to generate approximately 284 million 2×250 bp reads. Mate-pair reads were processed and used to scaffold contigs as described in the w2rap pipeline^12^ (https://github.com/bioinfologics/w2rap). Scaffolds less than 500 bp were removed from the final assembly. 245,820 scaffolds were generated comprising 4.11 Gb of sequence with an N50 of 86.6kb. Gaps comprise only 1.6% of the scaffolds (Table 1).

**Table 1:** Statistics for the different stages of the assembly process.

|  | **Contigs** | **Scaffolds** | **Dovetail** | **Hi-C** |
| --- | --- | --- | --- | --- |
| **N50** | 22.4kb | 86.67kb | 4.14Mb | / |
| **Number** | 786,696 | 245,820 | 128,283 | 8 |
| **Longest** | 352,153bp | 1,540,019bp | 22,832,123bp | 612,216,794bp |
| **Size** | 4.02Gb | 4.11Gb | 4.12Gb | 4.13Gb |

### Chromosome conformation capture

#### Dovetail

Golden Promise 10-day old leaf material was sent to Dovetail Genomics (Santa Cruz, CA, USA) for the construction of Chicago libraries. Dovetail extracted high molecular weight DNA and conducted the library preparations. The Chicago libraries were sequenced on an Illumina HiSeqX (Illumina, San Diego, CA, USA) with 150bp paired-end reads. Using the scaffold assembly as input, the HiRise scaffolding pipeline was used to build super scaffolds^13^.

#### Hi-C

The Hi-C library construction from one week old seedlings of Golden Promise was performed as per protocol described in Padmarasu *et al.*, 2019^14^ using DpnII for digestion of crosslinked chromatin. Sequencing of the Hi-C library was conducted on an Illumina HiSeq 2500 (Illumina, San Diego, CA, USA) with 101 bp paired-end reads. Super scaffolds from Dovetail were ordered and orientated to build the final pseudomolecule using the TRITEX assembly pipeline^10^, with a detailed user guide available (https://tritexassembly.bitbucket.io). This resulted in a final assembly of 4.13Gb and 7 chromosomes plus an extra chromosome containing the unassigned scaffolds.

### Completeness of the assembly

We used the spectra-cn function from the Kmer Analysis Toolkit (KAT)^15^ to check for content inclusion in the contigs and scaffolds. KAT generates a k-mer frequency distribution from the paired-end reads and identifies how many times k-mers from each part of the distribution appear in the assembly being compared. It is assumed that with high coverage of paired-end reads, every part of the underlying genome has been sampled. Ideally, an assembly should contain all k-mers found in the reads (not including k-mers arising from sequencing errors) and no k-mers not present in the reads.

The spectra-cn plot in Figure 1a generated from the contigs shows sequencing errors (k-mer multiplicity <20) appearing in black as these are not included in the assembly. The majority of the content appears in a single red peak indicating sequence that appears once in the assembly. The black region under the main peak is very small indicating that most of this content from the reads is present in the assembly. The content that appears to the right of the main peak and is present twice or three times in the assembly represents repeats.

**Figure 1:**
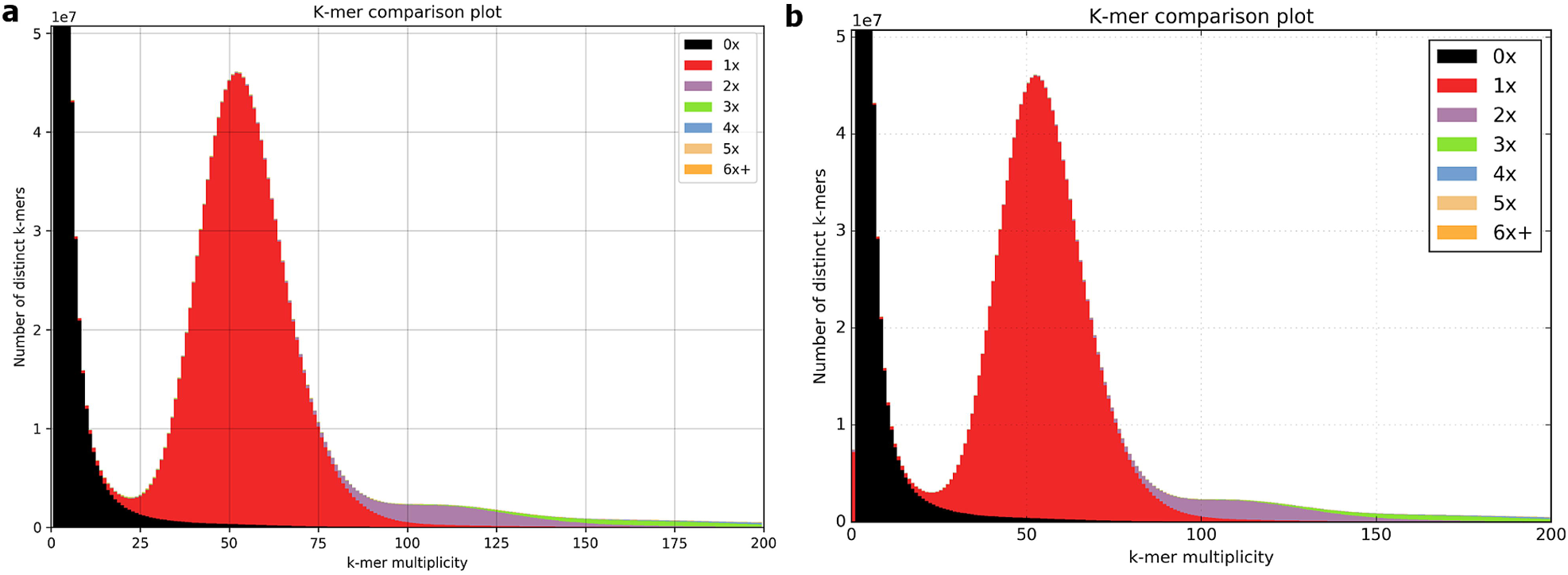
Spectra cn plots comparing k-mers from the paired-end reads to kmers in (a) the contig assembly and (b) the scaffold assembly

Scaffolds generally contain more miss-assemblies than contigs and this is reflected in the spectra-cn plot in Figure 1b generated from the scaffolds. The red bar at k-mer multiplicity 0 that is not present in the contigs spectra-cn plot reflects k-mers that appear in the scaffolds but do not appear in the reads. Approximately 7.2 million k-mers are represented in this region, less than 0.15% of the total.

### Repetitive Regions

The final assembly was analysed for repetitive regions using RepeatMasker (version 4.0.9)^16^ with the TREP Repeat library (trep-db_complete_Rel-16)^17^ and changing repetitive regions to lower case (-xsmall parameter) [repeat library downloaded from: http://botserv2.uzh.ch/kelldata/trep-db/downloadFiles.html]. The output of RepeatMask was condensed using the perl script “one-code-to-find-them-all”^18^ with the parameters --strict and --unknown. This identified 73.2% (2.95 Gb) of the Golden Promise assembly as transposable elements (Table 2) with almost all from the class of retroelements. The same analysis was also done for MorexV1 and MorexV2 showing that all three have very similar results (Table 2). Differences to the published results from MorexV1 and MorexV2 assembly^1,9,10^ are due to the different repeat libraries used.

**Table 2:** Identified repetitive elements in the Golden Promise assembly. Values represent percentage coverage of the genome.

|  | **Golden Promise** | **MorexV1** | **MorexV2** |
| --- | --- | --- | --- |
|  | 72.88 | 70.65 | 74.93 |
| **Class I: Retroelement** |  |  |  |
| LTR Retrotransposon | 63.16 | 62.25 | 64.25 |
| LTR/Copia | 19.87 | 21 | 20.94 |
| LTR/Gypsy | 42.97 | 40.93 | 42.99 |
| Unclassified LTR | 0.32 | 0.31 | 0.32 |
| **Non-LTR Retrotransposon** |  |  |  |
| LINE | 0.25 | 0.24 | 0.24 |
| SINE | 0.03 | 0.03 | 0.03 |
| **Class II: DNA Transposon** |  |  |  |
| DNA Transposon Superfamily | 8.25 | 7.39 | 8.97 |
| CACTA superfamily (DTC) | 7.77 | 6.92 | 8.49 |
| hAT superfamily (DTA) | 0.004 | 0.004 | 0.004 |
| Mutator superfamily (DTM) | 0.13 | 0.13 | 0.13 |
| Tc1/Mariner superfamily (DTT) | 0.2 | 0.19 | 0.2 |
| Harbinger superfamily (DTH) | 0.13 | 0.12 | 0.13 |
| Unclassified (DTX) | 0.02 | 0.02 | 0.02 |
| MITE (DXX) | 0.01 | 0.01 | 0.01 |
| Helitron (DHH) | 0.08 | 0.09 | 0.09 |
| Unclassified Element (XXX) | 0.46 | 0.3 | 0.74 |
| Simple Sequence Repeats | 0.63 | 0.36 | 0.59 |

## Data Validation and quality control

We used two approaches to evaluate the quality of the Golden Promise assembly based on gene content. The analysis was done for each of the steps along the assembly process. The first approach was done with BUSCO (Benchmarking Universal Single-Copy Orthologs, v3.0.2)^20,21^. It assesses the completeness of a genome by identifying conserved single-copy, orthologous genes. We used BUSCO with the plant dataset (embryophyta_odb9). For gene prediction BUSCO uses Augustus (Version 3.3)^22,23^. For the gene finding parameters in Augustus we set species to wheat and ran BUSCO in the genome mode (-m geno -sp wheat). Even the contig stage had already more complete single copy genes, 92.4%, in comparison to the published barley assembly from the cultivar MorexV1 with 91.5% (Figure 2a). Throughout the assembly process this improved to 95.2% of complete and single copy genes in the final pseudomolecule. This is very close to the recently published MorexV2 assembly with 97.2% of single copy genes. As expected, the number of fragmented sequences decreased during the assembly process from 2.8% of fragmented genes to only 1.1% in the pseudomolecule.

**Figure 2:**
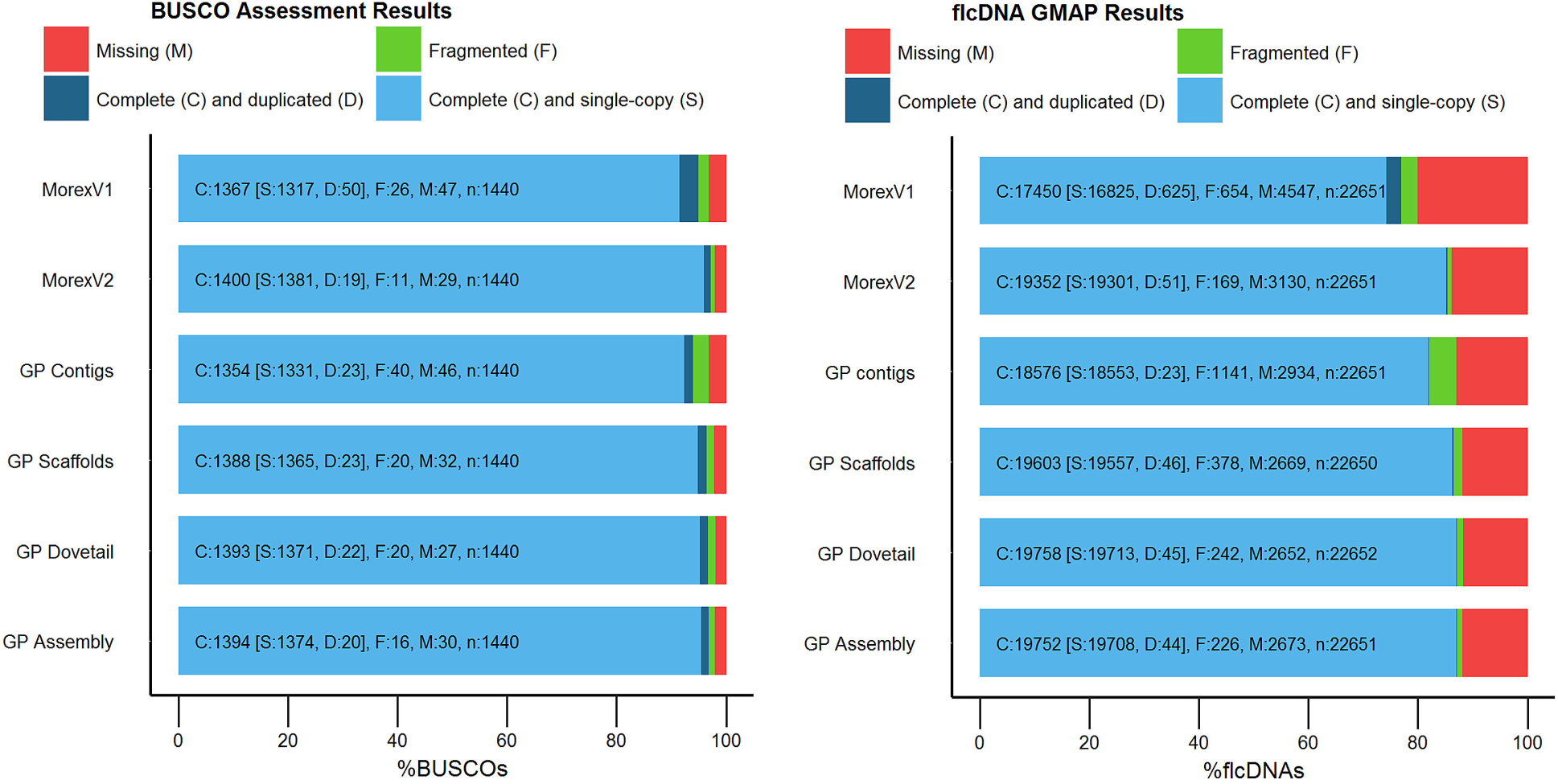
Completeness assessment of the Golden Promise assembly in comparison to the previous steps of the assembly process and the published barley reference MorexV1 for both the BUSCO analysis (a) and the flcDNA mapping analysis (b)

The second approach used a flcDNA dataset which consists of 22,651 sequences generated from the cultivar Haruna Nijo^24,25^. These sequences were created from 12 different conditions and representing a good snapshot of the barley transcriptome. They can be used to identify the number of retained sequences in the Golden Promise pseudomolecule and give an impression on the segmentation of the pseudomolecule, highlighted by cDNAs which have been split within or across chromosomes. The 22,651 flcDNAs were mapped to the Golden Promise pseudomolecule using Gmap (version 2018-03-25)^26^ with the following parameters: a minimum identity of 98% and a minimum trimmed coverage of 95%. The results for this dataset are very similar to the BUSCO analysis. The contigs already contained 81.4% of complete and single copy genes in comparison to the 73% of the MorexV1 reference (Figure 2b). The final assembly contained 87.1% of complete and single copy genes, 14% more than the barley reference MorexV1 and around 400 genes more in comparison to MorexV2 accounting for a difference of 1.9%. Similar to the BUSCO analysis the number of duplicated complete genes and the number of fragmented genes is decreased in the Golden Promise assembly. Again, the overall comparison to MorexV2 shows very similar results emphasising the high quality of both barley genomes.

## Conclusion

Here, we presented such an assembly that is an improvement on the currently available barley reference from the cultivar MorexV1^1,9^ and near-equivalent to the recently released MorexV2^10^. Importantly, it is a European 2-row cultivar, expanding barley genomic resources to European breeding material in contrast to the American 6-row cultivar Morex. We anticipate it will benefit the whole barley research community but will be especially useful for groups working on CRISPR-Cas9.

## Availability of supporting data

Raw reads have been deposited to the NCBI sequence read archive. Bioproject: PRJNA533066 [SRA: Paired end reads: SRR9291461, SRR9291462, SRR9291463, SRR9291464; Long mate pair reads: SRR9266823, SRR9266824, SRR9266825, SRR9266826, SRR9266827, SRR9266828; Dovetail reads: SRR9202370, SRR9202371, SRR9202372, SRR9202373, SRR9202374; Hi-C data: SRR8922888]

The reference assembly is available to download from figshare: 10.6084/m9.figshare.9332045. The reference assembly is available through the European Nucleotide Archive (ENA) under the following genome accession: GCA_902500625.

The reference sequence is also available to mine using an integrated gmap search. Transcript annotation is transferred from the BaRT transcriptome dataset^19^ and the TRITEX gene annotation^10^, using Gmap (version 2018-03-25) with the following parameters: -f 2 -n 1 --min-trimmed-coverage=0.8 --min-identity=0.9 (both files are available to download from figshare. BaRT: 10.6084/m9.figshare.9705278; TRITEX: 10.6084/m9.figshare.9705125).

Website for BLAST search access: https://ics.hutton.ac.uk/gmapper/

## List of abbreviations

BUSCO: Benchmarking Universal Single-Copy Orthologs
ENA: European Nucleotide Archive
LTR: Long terminal repeats
NCBI: National Centre for Biotechnology Information
PAM: protospacer-adjacent motif
sgRNA: single-guide RNA
TREP: TRansposable Elements Platform

## Consent for publication

Not applicable

## Competing interests

The authors declare no competing interests.

## Author contributions

RW designed and supervised the project. JW, DH, BC constructed the contigs and scaffolds and performed quality control. SP, AH and NS conducted the Hi-C experiment. MM performed pseudomolecule construction. LM developed the webpage. MS performed bioinformatics analysis and coordinated the project. MS, JW and RW wrote the manuscript with input from all other authors.

## Acknowledgments

The research leading to these results was funded by the H2020 European Research Council (ERC Shuffle, Project ID: 66918) to RW. We acknowledge the Scottish Government RESAS Strategic research program for supporting this research. MM and NS were supported by a grant from the German Federal Ministry of Education and Research (BMBF, FKZ 031B0190A ‘SHAPE’). We would like to thank and acknowledge Dovetail for their End-of-Year Matching Funds Grant to MS.

## References

1 International Barley Genome Sequencing Consortium. A physical, genetic and functional sequence assembly of the barley genome. Nature 491, 711–716, doi:10.1038/nature11543 (2012).

2 Mascher, M. et al. Genomic analysis of 6,000-year-old cultivated grain illuminates the domestication history of barley. Nat Genet 48, 1089–1093, doi:10.1038/ng.3611 (2016).

3 Hensel, G., Valkov, V., Middlefell-Williams, J. & Kumlehn, J. Efficient generation of transgenic barley: The way forward to modulate plant–microbe interactions. J Plant Physiol 165, 71–82, doi:10.1016/j.jplph.2007.06.015 (2008).

4 Ibrahim, A. S., El-Shihy, O. M. & Fahmy, A. H. Highly efficient Agrobacterium tumefaciens-mediated transformation of elite Egyptian barley cultivars. Am-Eurasian J Sustain Agric 4, 403–413 (2010).

5 Lim, W. L. et al. Method for hull-less barley transformation and manipulation of grain mixed-linkage beta-glucan. J Integr Plant Biol 60, 382–396, doi:10.1111/jipb.12625 (2018).

6 Murray, F., Brettell, R., Matthews, P., Bishop, D. & Jacobsen, J. Comparison of Agrobacterium-mediated transformation of four barley cultivars using the GFP and GUS reporter genes. Plant Cell Rep 22, 397–402, doi:10.1007/s00299-003-0704-8 (2004).

7 Belhaj, K., Chaparro-Garcia, A., Kamoun, S. & Nekrasov, V. Plant genome editing made easy: targeted mutagenesis in model and crop plants using the CRISPR/Cas system. Plant Methods 9, doi:10.1186/1746-4811-9-39 (2013).

8 Lawrenson, T. et al. Induction of targeted, heritable mutations in barley and Brassica oleracea using RNA-guided Cas9 nuclease. Genome Biol 16, 258, doi:10.1186/s13059-015-0826-7 (2015).

9 Mascher, M. et al. A chromosome conformation capture ordered sequence of the barley genome. Nature 544, 427–433, doi:10.1038/nature22043 (2017).

10 Monat, C. et al. TRITEX: chromosome-scale sequence assembly of Triticeae genomes with open-source tools. bioRxiv, 631648, doi:10.1101/631648 (2019).

11 Heavens, D., Accinelli, G. G., Clavijo, B. & Clark, M. D. A method to simultaneously construct up to 12 differently sized Illumina Nextera long mate pair libraries with reduced DNA input, time, and cost. BioTechniques 59, 42–45, doi:10.2144/000114310 (2015).

12 Clavijo, B. et al. W2RAP: a pipeline for high quality, robust assemblies of large complex genomes from short read data. bioRxiv, 110999, doi:10.1101/110999 (2017).

13 Putnam, N. H. et al. Chromosome-scale shotgun assembly using an in vitro method for long-range linkage. Genome Res 26, 342–350, doi:10.1101/gr.193474.115 (2016).

14 Padmarasu, S., Himmelbach, A., Mascher, M. & Stein, N. in Plant Long Non-Coding RNAs: Methods and Protocols (eds Julia A. Chekanova & Hsiao-Lin V. Wang) 441–472 (Springer New York, 2019).

15 Mapleson, D., Venturini, L., Kaithakottil, G. & Swarbreck, D. Efficient and accurate detection of splice junctions from RNAseq with Portcullis. bioRxiv, 217620, doi:10.1101/217620 (2017).

16 Smit, A., Hubley, R. & Green, P. RepeatMasker Open-4.0, <http://www.repeatmasker.org> (2013-2015).

17 Wicker, T., Matthews, D. E. & Keller, B. TREP: a database for Triticeae repetitive elements. Trends Plant Sci 7, 561–562, doi:10.1016/S1360-1385(02)02372-5 (2002).

18 Bailly-Bechet, M., Haudry, A. & Lerat, E. “One code to find them all”: a perl tool to conveniently parse RepeatMasker output files. Mobile DNA 5, 13, doi:10.1186/1759-8753-5-13 (2014).

19 Rapazote-Flores, P. et al. BaRTv1.0: an improved barley reference transcript dataset to determine accurate changes in the barley transcriptome using RNA-seq. bioRxiv, 638106, doi:10.1101/638106 (2019).

20 Simao, F. A., Waterhouse, R. M., Ioannidis, P., Kriventseva, E. V. & Zdobnov, E. M. BUSCO: assessing genome assembly and annotation completeness with single-copy orthologs. Bioinformatics 31, 3210–3212, doi:10.1093/bioinformatics/btv351 (2015).

21 Waterhouse, R. M. et al. BUSCO Applications from Quality Assessments to Gene Prediction and Phylogenomics. Mol Biol Evol 35, 543–548, doi:10.1093/molbev/msx319 (2018).

22 König, S., Romoth, L. W., Gerischer, L. & Stanke, M. Simultaneous gene finding in multiple genomes. Bioinformatics 32, 3388–3395, doi:10.1093/bioinformatics/btw494 (2016).

23 Stanke, M., Steinkamp, R., Waack, S. & Morgenstern, B. AUGUSTUS: a web server for gene finding in eukaryotes. Nucleic Acids Res 32, W309–W312, doi:10.1093/nar/gkh379 (2004).

24 Matsumoto, T. et al. Comprehensive Sequence Analysis of 24,783 Barley Full-Length cDNAs Derived from 12 Clone Libraries. Plant Physiol 156, 20–28, doi:10.1104/pp.110.171579 (2011).

25 Sato, K. et al. Development of 5006 Full-Length CDNAs in Barley: A Tool for Accessing Cereal Genomics Resources. DNA Res 16, 81–89, doi:10.1093/dnares/dsn034 (2009).

26 Wu, T. D. & Watanabe, C. K. GMAP: a genomic mapping and alignment program for mRNA and EST sequences. Bioinformatics 21, 1859–1875, doi:10.1093/bioinformatics/bti310 (2005).

